# ResiRuler: A Toolkit for Visualizing Residue-Residue Distances and Structural Changes in Biomolecular Models

**DOI:** 10.64898/2026.08.19.745761

**Authors:** Timothy H. Baker, Melanie D. Ohi, Wilhelm Salmen

**Affiliations:** Life Sciences Institute, University of Michigan, Ann Arbor, MI; Department of Cell and Developmental Biology, University of Michigan, Ann Arbor, MI

**Keywords:** protein structure comparison, conformational variability, residue-level analysis, structural ensembles, distance difference mapping, molecular visualization, macromolecular complexes, Cartesian coordinate analysis, ChimeraX, PyMOL

## Abstract

Proteins and their associated complexes often adopt multiple conformations, with the transitions between these states playing a critical role in biological function. However, the resulting structural heterogeneity can be challenging to visualize and communicate, often requiring manual inspection and time-consuming annotation of biomolecular structures. To address this, we developed ResiRuler, a local, browser-based tool that uses inter-residue distance measurements to quickly quantify atomic displacements and map changes in internal geometry across ensembles of related protein structures. By converting structural differences into residue-pair distance changes, ResiRuler enables rapid identification of regions undergoing coordinated motion, local rearrangement, or large-scale conformational change. The resulting visualizations can be exported as scripts for PyMOL and ChimeraX, allowing users to explore conformational differences and generate publication-quality molecular figures in their preferred visualization environment. Using atomic models in Macromolecular Crystallographic Information File (mmCIF) file format, ResiRuler aligns multiple structures and measures structural variation across models facilitating visualization and presentation of these differences. This allows for rapid visualization of which regions of proteins change among ensembles of structures. The program is available for download at https://github.com/tbaker67/ResiRuler on macOS and Linux operating systems.

**Broad Audience Statement:** Proteins and molecular machines often change shape to perform their biological functions, but comparing these movements across structural models can be slow and difficult. ResiRuler makes this process easier by measuring and visualizing residue-level structural changes through an accessible browser-based tool. By helping researchers quickly identify coordinated motions and local rearrangements, ResiRuler can improve interpretation of protein structures and support clearer communication of molecular mechanisms.

## Introduction

Recent advances in structural biology techniques, including nuclear magnetic resonance spectroscopy (NMR), X-ray crystallography, and cryo-electron microscopy (cryo-EM), have made it easier to determine protein structures with increasing resolution and scale. In parallel with the growing number of experimental structures, deep learning–based methods such as AlphaFold and ESMFold have revolutionized structural biology by enabling increasingly accurate predictions of individual proteins and protein assemblies, further increasing the structural landscape available for analyses^1–6^. As a result, the number and diversity of atomic models have expanded, with many proteins and complexes now represented by multiple structures that capture distinct conformational and functional states ^7^. Reflecting this growth, the Protein Data Bank (PDB) currently contains more than 227,000 experimentally determined structures and over 1,000,000 Computed Structural Models (CSMs) ^8^.

This rapid growth in structural data has also increased the complexity of comparative protein analysis. Many newly available structures represent multi-chain assemblies, large macromolecular complexes, or alternative conformational states rather than isolated monomeric proteins. These structures create new opportunities to investigate variability across related models, but they also present challenges for quantifying and visualizing differences in a manner that is both scalable and interpretable. As structural datasets continue to grow, there is a pressing need for robust tools capable of comparing local and global structural variation across multiple models while producing clear, publication-ready visualizations. While these advances have expanded the structural data available for systematic analyses of protein structure, dynamics, and function across conformational landscapes and among evolutionary related proteins, approaches for rapidly comparing and visualizing multiple structures have not kept pace.

One common approach for visualizing structural differences is molecular morphing, where one model is gradually transformed into another using a continuous trajectory, providing an intuitive representation of conformational change^9,10^. However, morphing is primarily qualitative and does not directly quantify local structural shifts. For more direct comparisons between models, researchers often rely on metrics such as root-mean-square deviation (RMSD) or other distance-based measures to summarize structural similarity between two models. Although useful, these metrics reduce structural differences to a single global value, obscuring localized conformational changes and limiting interpretation of the specific regions that differ between models.

Localized structural comparisons provide a complementary approach by measuring differences at the residue or atom level. For example, atom displacement plots and distance difference maps can reveal the location and magnitude of structural divergence between models. Tools such as FATCAT 2.0 support interactive visualization of these localized differences and illustrate the utility of residue-level structural metrics ^11^. However, many existing implementations are designed primarily for interactive inspection rather than publication-quality figure generation, and they are not optimized for comparing large datasets or complex multi-chain assemblies.

To address these limitations, we introduce ResiRuler, a Python-based web tool for quantifying and visualizing structural differences between two or more atomic models at the residue level. ResiRuler compares Cartesian coordinate-derived distances at the c_a_- backbone or amino acid side-chain level of atomic models and maps these differences directly onto protein structures. This enables users to visualize and quantify variability across structural ensembles, related homologs, predicted models, or experimentally determined conformational states. ResiRuler is designed for stand-alone exploratory analysis through an interactive web interface and can also export high-quality visualization scripts compatible with ChimeraX^9^ and PyMOL^12^. Together, these features provide a program with an accessible workflow for analyzing local structural variation in increasingly large and complex protein structure datasets.

## Results

### Software Overview

ResiRuler is a locally run software tool that uses Streamlit^13^, an open source Python framework, to provide a browser-based interface for residue-level comparison of macromolecular structures. Through this interface, users can upload structural models, align them to either a designated reference model or map corresponding residues across models then calculate residue-level structural metrics, and generate interactive visualizations for comparing multiple models without requiring command-line interaction (Figure 1). The only required inputs are three-dimensional (3D) atomic coordinate files for two or more structures to be compared. ResiRuler accepts coordinate files in Macromolecular Crystallographic Information File (mmCIF) format^14,15^, providing compatibility with atomic models derived from NMR, X-ray crystallography, or cryo-EM data, as well as structures generated by tools such as AlphaFold.

**Figure 1.**
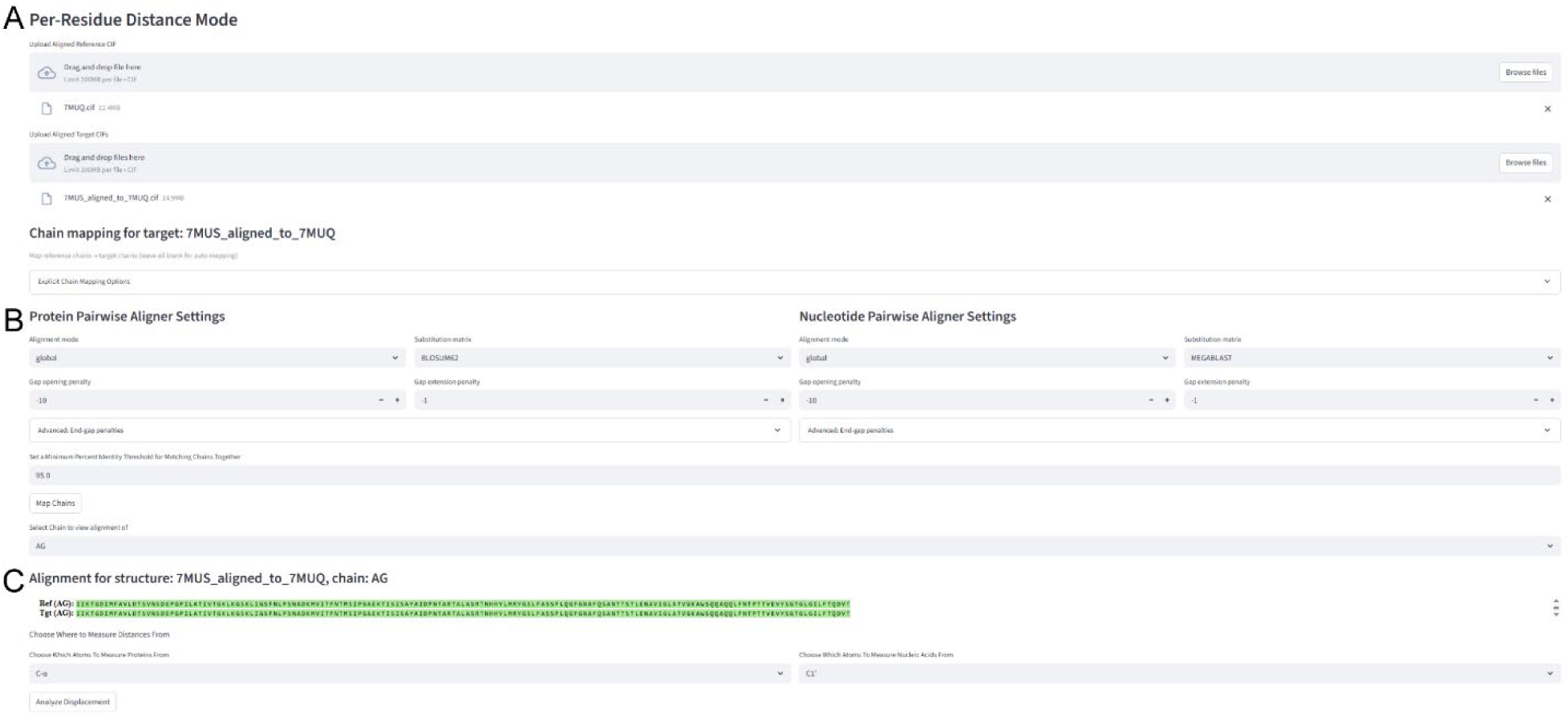
ResiRuler interface shows default settings for uploading and comparing structures. (A) Aligned reference and comparison models can be uploaded as mmCIF files. (B) Protein pairwise aligner settings allow users to adjust alignment parameters for insertions, deletions, and substitutions, as well as manually assign chain pairs when needed. (C)The chain-alignment panel shows chain assignments, with green text indicating accepted matches. For residue-level comparisons, users can select Cα atoms, Cβ atoms, or side-chain centroids before computing pairwise distance differences or per- residue displacements.

The overall ResiRuler workflow is summarized in Figure 2A. Users first upload a reference structure and one or more additional structures into the designated fields of the interface. Because distance-based residue comparisons require all the structures to share a common spatial frame, the additional structures must be aligned to the reference before analysis. Users may either upload previously aligned mmCIF files or use the built-in alignment module in ResiRuler (Figure 1A). When automatic alignment is selected, ResiRuler applies TM-align^16^, an algorithm for sequence independent protein structure alignment, to perform rigid-body superposition optimized by TM-score. The interface also allows users to remove residues that lack a corresponding match in the reference structure, reducing computational cost and focusing downstream analyses to only the structurally comparable regions .

**Figure 2.**
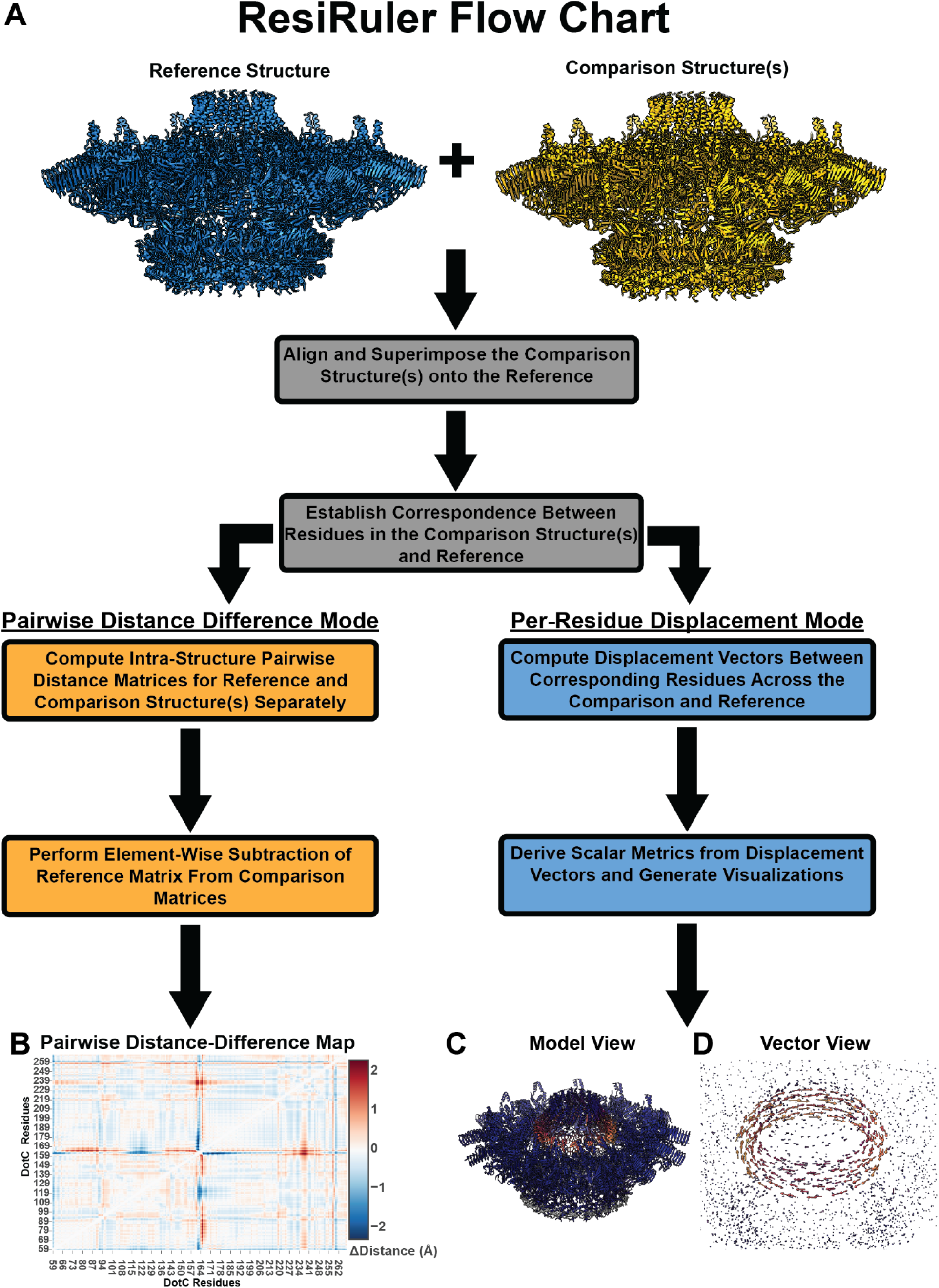
ResiRuler workflow from model input to visualization and export of residue-level comparison outputs. (A) Users provide a reference model and one or more target models, which may be prealigned or aligned automatically within ResiRuler. Users then select either pairwise distance-difference mode, shown in orange, or per-residue displacement mode, shown in blue. In both modes, ResiRuler first establishes chain and residue correspondences between the reference and target structures and displays the selected visualization in built-in structure viewers. (B) In pairwise distance-difference mode, ResiRuler calculates pairwise residue-distance matrices for the reference and target structures and subtracts the reference matrix from the target matrix to generate distance-difference maps and plots showing changes in distance. Red indicates residue pairs that are farther apart in the target model than in the reference model, whereas blue indicates residue pairs that are closer together in the target model than in the reference model. Distance differences are reported in angstroms (Å). (C-D) In per-residue displacement mode, the software calculates displacement vectors between corresponding residues and derives distance-based metrics used to generate visualization scripts that (C) color-code structural models by displacement magnitude and (D) display displacement vectors. For both displacement and pairwise distance-difference modes, exported outputs are bundled into a results folder containing visualization scripts, visualization-ready models, and associated comma-separated values files.

After spatial alignment, ResiRuler establishes chain and residue correspondences between the reference and each structure being compared using pairwise sequence alignment implemented through Biopython sequence aligners^17^. By default, ResiRuler determines residue correspondences uses the Needleman–Wunsch algorithm with a BLOSUM62 substitution matrix, a gap-opening penalty of −10, and a gap-extension penalty of −1^18^. These parameters can be adjusted to accommodate differences in sequence similarity, insertions, deletions, or structural completeness (Figure 1B). For each chain, ResiRuler generates residue-level alignment views that allow users to inspect residue correspondences, identify gaps, and verify that unresolved or unmatched regions are handled appropriately (Figure 1C).

Following the establishment of residue correspondence among the structures, users select the chains to include in the analysis and define visualization color schemes. ResiRuler then computes residue-level distance metrics using Cα atoms, Cβ atoms, or side-chain centroids (Figure 1C). These multiple options allow users to assess different aspects of structural variation, from backbone displacement to side-chain-level rearrangements. Results are displayed in integrated molecular viewers, enabling immediate visual inspection of conformational similarities and differences across the structural ensemble.

ResiRuler exports a structured results directory containing the aligned models used for analysis, comma-separated values (CSV) files of computed metrics, and visualization scripts compatible with viewing in PyMOL and ChimeraX^9,12^. These outputs allow users to further examine structural differences, customize figures, and generate publication-quality structural representations. ResiRuler has two distinct but related visualization modes: **Pairwise Distance Difference**, which quantifies changes in inter-residue distances, and **Per-Residue Displacement**, which visualizes residue-level positional shifts relative to the reference structure (Figure 2A).

The **Pairwise Distance Difference Mode** identifies local and global conformational differences by first calculating intra-structure pairwise distance matrices for the reference and the aligned comparison structure. Both matrices are constructed from shared residue correspondences across the structures, allowing elements to be directly compared across structures. Because of this, subtracting the reference matrix from a target matrix produces a distance-difference matrix that reports how inter-residue distances change between models.

These distance-difference matrices can be displayed as heat maps or summarized as residue-level metrics, such as the average distance change per residue (Figure 2B). By applying a distance threshold to the pairwise matrices, users can also generate contact maps that highlight residue pairs within a user-defined cutoff, measured in Å. Mapping gained and lost residue contacts between structures highlights regions where conformational changes may alter domain interfaces, binding surfaces, or other protein-protein interactions.

The **Per-Residue Displacement Mode** focuses on residue-level positional changes across aligned structures. For each corresponding residue, ResiRuler calculates a displacement vector from the coordinate difference between the residue in reference structure and its counterpart in the comparison structure. The magnitude of this vector reports the extent of residue displacement, and its direction indicates the orientation of that displacement in 3D space.

Residue displacements can be visualized on the reference structure as color gradients indicating magnitude or as arrows indicating both displacement magnitude and direction (Figure 2C,D). To improve visual clarity, users can adjust the number of displacement vectors shown in the interface display as well as color scaling. Examining both the magnitude and direction of residue-level displacements allows users to identify flexible domains, hinge-like motions, and the overall geometry of conformational change between models.

### Example application: structural comparison of the *Legionella pneumophila* Dot/Icm type IV secretion system (Dot/Icm T4SS)

ResiRuler is well suited for comparing large, multi-chain assemblies, particularly those with symmetry mismatches, conformational variability, and/or incomplete one-to-one chain correspondence. As an example, we applied ResiRuler to structures of the *L. pneumophila* Dot/Icm T4SS. The Dot/Icm T4SS is a large inner and outer membrane spanning molecular machine that translocates effectors from *L. pneumophila* into host cells^19,20^.

The outer-membrane core complex (OMCC), composed of at least nine proteins, is the part of the T4SS localized in the periplasm, the space between the bacterial outer and inner membrane. The OMCC is organized into four sub-regions: a 16-fold symmetric dome, a 13-fold symmetric outer membrane cap (OMC), an 18-fold symmetric periplasmic ring (PR), and a 5-fold symmetric stalk^19–21^. The single particle cryo-EM structure of the OMCC included the dome, OMC, and PR and resolved 158 protein chains that mapped to: 31 DotF, 26 DotD, 18 DotG, 18 DotH, and 13 copies each of DotC, DotK, Dis1, Dis2, and Dis3^21^. 3D variability analysis (3DVA) further showed that the OMCC samples distinct conformational states, especially in the spatial orientation between the dome, OMC, and PR regions^21^. The combination of extensive chain multiplicity, symmetry mismatches, and conformational variation makes this complex an ideal test case for ResiRuler.

In the 3DVA-derived conformation states, the most prominent differences involve shifts in the position of DotG relative to the OMC and PR^21^. However, these changes are difficult to easily visualize and quantify because DotG spans multiple symmetry-mismatched regions of the OMCC. There are 18 DotG protomers in the PR, 13 domains of DotG extend from the PR into the OMC, and 16 DotG a-helices form the dome^21^. In addition, the dome, OMC, and PR structures were each refined using applied symmetry, eliminating the physical connections across these symmetry-mismatched regions and preventing DotG from being traced continuously through the OMCC. This makes the 3DVA states of the Dot/Icm T4SS difficult to compare by visual inspection alone and highlights the need for residue-level approaches to quantify and visualize conformational differences among structures.

To address this, we used ResiRuler to analyze and visualize conformational differences among the 3DVA-derived models (PDB: 7MUQ, 7MUS, 7MUY, 7MUW, and 7MUV) (Figure 3A,C; Supplemental Figure 1)^21^. For all analyses the model generated from 3DVA map 1 served as the reference structure (PDB: 7MUQ). In Per-Residue Displacement Mode, residue displacements between the reference and comparison structures were quantified at the residue level and visualized as vectors (Figure 3D) and color scaling directly on the reference model for each aligned comparison structure (Figure 3C and Supplemental Figure 1). These vectors indicate the magnitude and direction of residue-level positional shifts. In Pairwise Distance Difference Mode, pairwise residue distance-difference matrices between the reference and target structures are calculated and displayed as heat maps (Figure 4).

**Figure 3.**
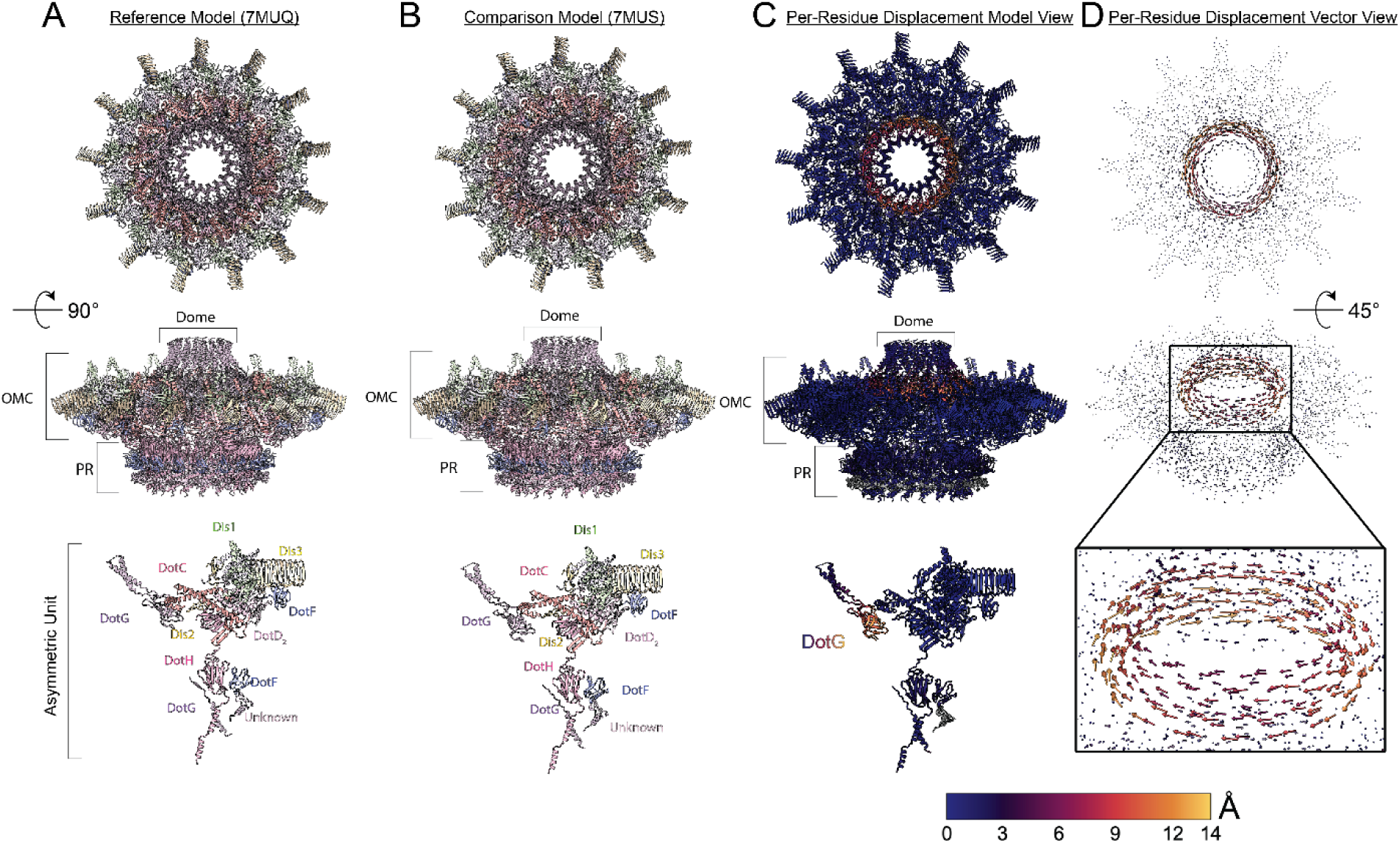
ResiRuler Per-Residue Displacement Mode allows for visualization of Cα positional differences between two 3DVA-derived models of the Dot/Icm T4SS. (A- B) Structures of 3DVA-derived models of the DotIcm T4SS. 7MUQ is the reference and 7MUS is the comparison structure. (C) Reference structure (7MUQ) colored by residue- level distance differences calculated by ResiRuler in Per-Residue Displacement Mode. (A-C) Top panels: *en face* view (inner- to outer- membrane view); Middle panels: Side view (90° rotation from top panels). Dome, OMC, PR are labeled; Bottom panels: Asymmetric unit. Position of individual T4SS components are colored and labeled in the asymmetric unit. (D) Vector representation of Cα displacement from residues in 7MUQ to their corresponding residues in 7MUS. Arrows indicate 3D direction of differences between the two structures. Colors correspond to residue differences spanning 0-14 Å as shown in the bar. Top panel: *en face* view (inner- to outer- membrane view); Middle panels: Side view (45° rotation from top panel); Bottom panel: enlargement of boxed region in middle panel.

**Figure 4.**
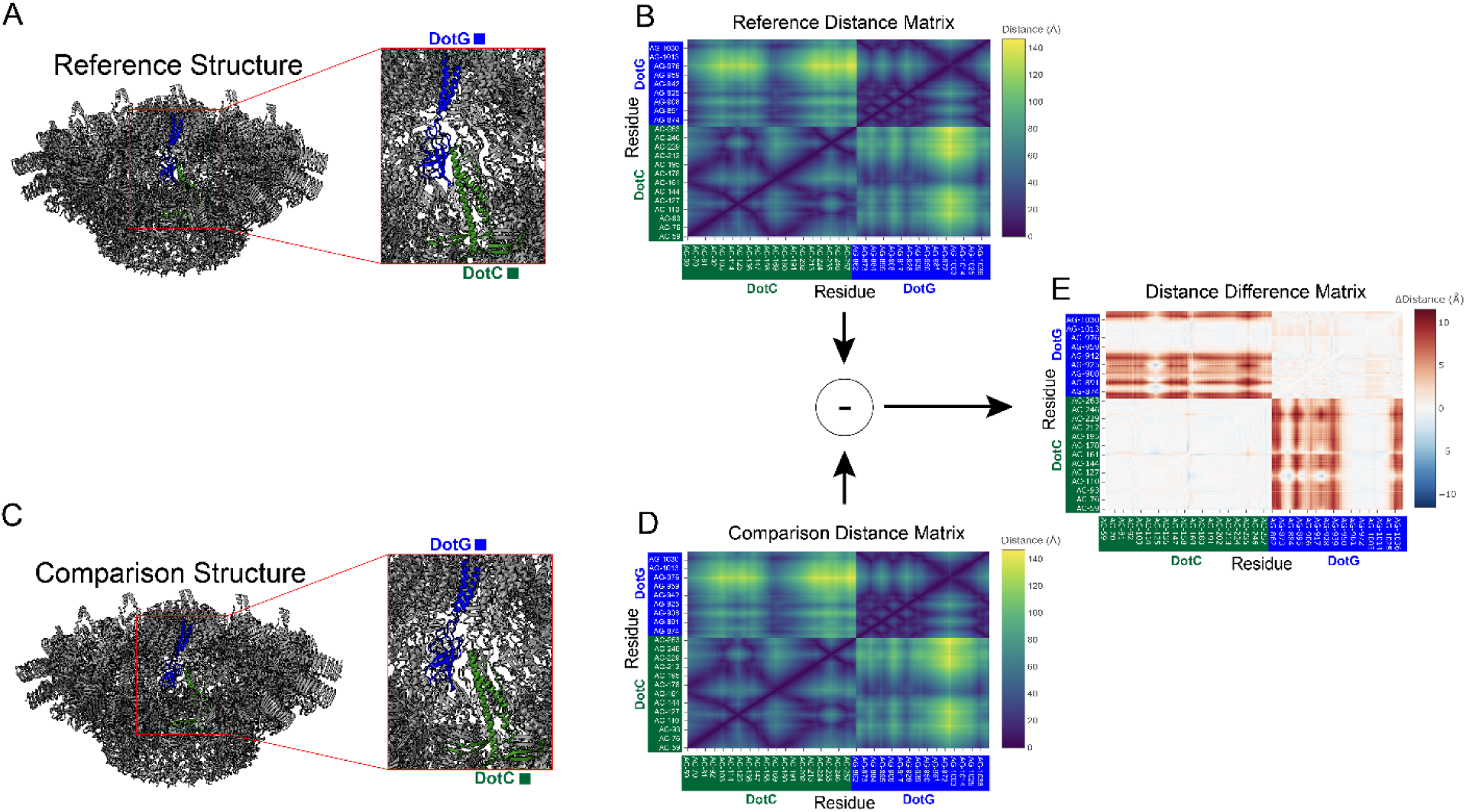
Distance-difference matrices reveal residue-pair structural shifts. ResiRuler compares reference and target structures using shared residue correspondence. In this example, a subunit of DotC, shown in green, and DotG, shown in blue, were selected for comparison. (A,C) ResiRuler calculates all pairwise residue distances within and between DotC and DotG for both the reference (7MUQ) and target structure (7MUS). The reference distance matrix (B) is then subtracted from the target distance matrix (D) to generate a distance-difference matrix in which each element represents the change in distance between a residue pair in the target structure relative to the reference. In the resulting distance-difference plot (E), red indicates increased distance between a residue pair, whereas blue indicates decreased distance.

These analyses identified the orientation of DotG within the dome, relative to the OMC and PR, as the primary conformational difference among these 3DVA-derived models. Comparison of 7MUQ with 7MUS revealed one of the largest changes between the models, with an approximately 15 Å shift of the DotG helix within the dome region accompanied by radial rotation relative to the OMC and PR (Figures 3C,D). This observation is corroborated when compared using Pairwise distance difference where we observe Pairwise residue differences of up to 10 Å between residues within DotG and DotC (Figure 4E). Comparisons of the other 3DVA-derived models against the 7MUQ reference structure showed additional DotG displacements, including changes in the positions of the membrane-facing helices at the tip of the dome (Supplemental Figure 1B,C,D) and an ∼6 Å shift of the dome relative to the PR (Supplemental Figure 1E). These ResiRuler visualizations show how residue-level displacements across multiple maps correspond to larger domain movements within the Dot/Icm T4SS. This intuitive approach for visualizing conformational changes across large structural ensembles facilitates the development of new hypotheses about how these changes may contribute to substrate accommodation and transport.

## Discussion

ResiRuler provides a practical framework for residue-level comparison of related macromolecular structures. By combining structure alignment, residue correspondence mapping, quantitative distance measurements, and interactive visualization in a browser-based interface, the software lowers technical barriers for identifying and mapping conformational differences across ensembles of experimentally determined structures or predicted models.

The two visualization modes provide complementary views of structural variation. “Pairwise Distance Difference Mode” evaluates changes in inter-residue distances, enabling identification of coordinated structural rearrangements, domain expansion or contraction, and changes in internal structural relationships that may not be apparent from global superposition alone. “Per-Residue Displacement Mode” evaluates direct residue- level positional shifts relative to a reference structure, providing an intuitive representation of local movement and its directionality. Together, these measurements allow conformational variation among structures to be examined from both internal-distance and coordinate-displacement perspectives.

ResiRuler is especially useful for comparing structural ensembles of multi-chain assemblies and systems that are symmetric or pseudo-symmetric, where manual chain and residue matching can be tedious. Automated chain assignment and residue mapping enable comparisons among structurally related models while preserving user oversight of residue correspondences. The ability for the user to exclude unmatched residues helps ensure that calculated metrics reflect comparable structural regions rather than sequence gaps, insertions, or unresolved density.

The exported output also supports visualization in molecular graphics programs commonly used by structural biologists, biochemists, and bioinformaticians. ResiRuler generates aligned coordinate files, CSV files containing computed residue-level metrics, and visualization scripts for PyMOL and ChimeraX^9,12^. These outputs allow users to reproduce analyses, customize structural representations, and generate publication-quality figures using established molecular visualization programs.

Several limitations should be considered when interpreting ResiRuler results. The accuracy of residue-level comparisons depends on the quality of the initial structural alignment and residue mapping. Highly divergent structures, extensive insertions or deletions, incomplete models, or ambiguous chain relationships may require manual inspection or explicit chain assignment. In addition, displacement values are reference-dependent and may reflect alignment choices rather than biologically meaningful conformational changes. As with other coordinate-based approaches, residue-level differences should be interpreted in the context of model quality, structural uncertainty, and the biological question being addressed.

Future development could include support for additional coordinate formats, expanded ensemble-level statistical analyses, improved treatment of symmetric chain ambiguity, and integration of model-confidence or structure-quality metrics. In its current form, ResiRuler offers a streamlined and reproducible platform for residue-level structural comparison, helping users quantify, visualize, and communicate conformational differences across related macromolecular models. Thus, although additional features could further expand its utility, ResiRuler already provides a powerful and accessible tool for analyzing structural ensembles and interpreting conformational heterogeneity.

## Methods

### Automatic Chain Alignment Algorithm

For automated chain alignment, ResiRuler performs pairwise amino acid sequence alignments between chains in the reference and target structures. Pairwise alignments are calculated using the Needleman–Wunsch algorithm as implemented in the Bio.Align package from Biopython^17,18^. By default, alignments use the BLOSUM62 substitution matrix, a gap-opening penalty of −10, and a gap-extension penalty of −1, although these parameters can be modified by the user^22^ (Figure 1B).

For each possible reference–target chain pairing, ResiRuler records the sequence alignment score and calculates a RMSD using the aligned residue positions. These two quantities are combined into a single cost function used for chain assignment:

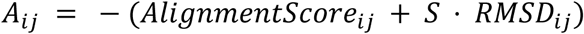

Where *i* and *jj* denote chains in the reference and target structures, respectively, and *S* is a scaling factor applied to the RMSD term. By default, the formulation favors chain pairs with high sequence alignment scores while allowing RMSD to serve as a secondary criterion, particularly for distinguishing identical or near-identical chains in symmetric or pseudo-symmetric assemblies. However, the user is also able to adjust the scaling factor S to increase the influence of structural similarity in chain assignments, which may be preferable when comparing distantly related sequences or assemblies where sequence identity alone is insufficient to resolve chain correspondences

If the sequence identity between two chains falls below a user-defined threshold, that pairing is assigned an artificially high cost. This prevents low-confidence pairings from being selected during assignment and allows unmatched or poorly matched chains to remain unpaired.

All pairwise chain costs are assembled into a cost matrix and passed to the linear_sum_assignment function from the scipy.optimize package^23^. This function implements a modified Jonker–Volgenant algorithm to identify the optimal one-to-one assignment that minimizes the total chain-pairing cost^24^. Pairings with artificially high costs are subsequently discarded, producing the final set of accepted chain correspondences.

## Global Residue Mapping

After chain correspondences have been established, either automatically or through user-specified mappings, ResiRuler constructs a global residue mapping across the structural ensemble. For each reference chain that is successfully matched to a corresponding chain in every target structure, ResiRuler iterates through the residues in the reference chain and identifies the corresponding residues in each target based on the pairwise sequence alignments.

A reference residue is included in the global mapping only when a valid corresponding residue is present in all selected target structures. These matched residues are stored in parallel arrays, with one array for each structure. Residues occupying the same index across the arrays represent equivalent residue positions across the reference and target structures. This data structure enables consistent and efficient residue-level comparisons across the ensemble.

Residues lacking valid correspondences, including gaps, unresolved positions, or unmatched insertions, are excluded from the global mapping. This ensures that downstream distance and displacement calculations are performed only on structurally comparable residue positions.

## Pairwise Distance Difference Mode

Pairwise Distance Difference Mode visualization is generated from the globally mapped residue-coordinate arrays. For each structure, ResiRuler calculates a pairwise residue distance matrix by computing the Euclidean distance between all globally aligned residue positions. Depending on the user-selected analysis mode, residue positions may be represented by Cα atoms, Cβ atoms, or side-chain centroids (Figure 1C).

Because all distance matrices are constructed from the same global residue mapping, corresponding matrix elements represent the same residue pairs across all structures. This enables direct element-wise comparison between the reference and target matrices. For a given target structure, the distance-difference matrix is calculated as:

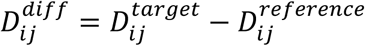

Where 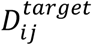 is the distance between residues *i* and *jj* in the target structure, and 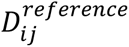 is the corresponding distance in the reference structure. Positive values indicate residue pairs that are farther apart in the target structure than in the reference, whereas negative values indicate residue pairs that are closer together.

Distance-difference matrices can be displayed as heat maps to visualize pairwise residue-distance changes across the structure (Figure 2B). ResiRuler also summarizes these pairwise differences into residue-level metrics, such as the average absolute distance change associated with each residue.

## Per-Residue Displacement Mode

Per-Residue Displacement Mode visualization uses the same globally mapped residue-coordinate arrays generated during residue mapping (Figure 2A). For each residue, ResiRuler calculates a displacement vector describing the positional difference between the reference coordinate and the corresponding coordinate in a target structure:

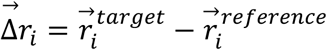

where 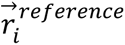 and 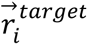 are the coordinates of residue *i* in the reference and target structures, respectively. The magnitude of this vector is calculated as:

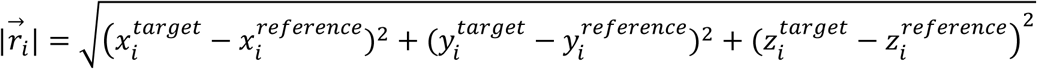

The vector magnitude reports the extent of residue-level displacement, whereas the vector direction indicates the orientation of movement from the reference structure to the target structure. As in Pairwise Distance Difference Mode, residue coordinates can be represented using Cα atoms, Cβ atoms, or side-chain centroids (Figure 1C).

Displacement magnitudes are normalized and mapped to a color gradient using Matplotlib colormaps^25^ (Figure 2C). The color scale may be determined automatically from the minimum and maximum displacement values or specified manually by the user. Displacement vectors are visualized in three dimensions using Plotly, with each vector originating at the reference residue coordinate and pointing toward the corresponding residue in the target structure^26^ (Figure 2D). To improve readability in large structures, users can adjust the number of displayed vectors using the interface. Additionally, the resulting values can be displayed directly on the reference model or exported for visualization in external molecular graphics programs.

## Exported Script Generation

To support downstream visualization and figure generation, ResiRuler exports scripts and data files compatible with ChimeraX and PyMOL^9,12^.

For ChimeraX, ResiRuler generates .defattr, .bild, and command-script files. The .defattr files encode residue-level attributes that can be applied to molecular models for color mapping. The .bild files define graphical objects, including arrows and lines used to represent displacement vectors. ChimeraX command scripts automate model loading, application of residue attributes, rendering of vector objects, and display configuration. These files allow users to reproduce ResiRuler visualizations directly in ChimeraX.

For PyMOL, ResiRuler generates .pml scripts that automate structure loading, visual styling, residue-level coloring, and display configuration. These scripts allow users to further customize structural representations, generate publication-quality images, and integrate ResiRuler outputs into existing PyMOL-based workflows.

In addition to visualization scripts, ResiRuler exports aligned coordinate files and comma-separated values files containing all calculated residue-level metrics. These files provide a reproducible record of the analysis and allow users to perform additional statistical analysis or visualization outside the ResiRuler interface.

## Data Availability

The program is available for download at https://github.com/tbaker67/ResiRuler on macOS and Linux operating systems

## Acknowledgements

We are grateful to Arwen E. Frick-Cheng for reviewing the manuscript, Chia-Yu Kang and Sid Ramesh for providing test models, and all members of the Melanie D. Ohi and Herman Fung laboratories for their valuable feedback on the software concepts.

---

**Supplemental Figure 1.**
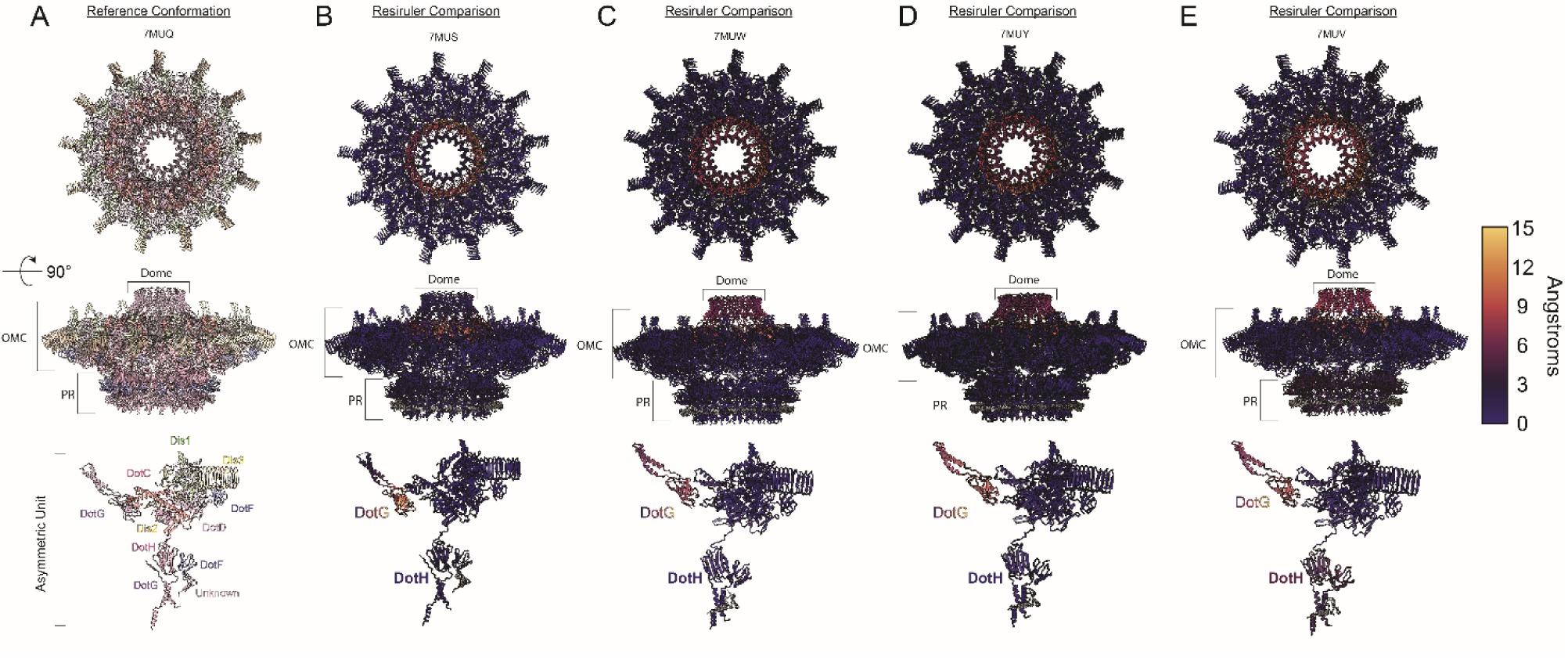
R**e**siRuler **structure-comparison using Per-Residue Displacement Mode against multiple target model outputs.** Structures of 3DVA- derived models of the Dot/Icm T4SS. 7MUQ (A) is the reference structure colored by subunit. 7MUS (B), 7MUW (C), 7MUY (D), and 7MUV (E) are the comparison structures, colored by displacement magnitude. Top panels: *en face* view (inner- to outer- membrane view); Middle panels: Side view (90° rotation from top panels). Dome, OMC, PR are labeled; Bottom panels: Asymmetric unit.

